# A two-way molecular dialogue between embryo and endosperm required for seed development

**DOI:** 10.1101/809368

**Authors:** N. M. Doll, S. Royek, S. Fujita, S. Okuda, A. Stintzi, T. Widiez, M. Hothorn, A. Schaller, N. Geldner, G. Ingram

**Affiliations:** Laboratoire Reproduction et Développement des Plantes, University of Lyon, ENS de Lyon, UCB Lyon 1, CNRS, INRA, F-69342, Lyon, France; Department of Plant Physiology and Biochemistry, University of Hohenheim, Stuttgart, Germany; Department of Plant Molecular Biology, University of Lausanne, 1015 Lausanne, Switzerland; Structural Plant Biology Laboratory, Department of Botany and Plant Biology, University of Geneva, 1211 Geneva, Switzerland

## Abstract

The plant embryonic cuticle is a hydrophobic barrier deposited *de novo* by the embryo during seed development. At germination it protects the seedling from water loss and is thus critical for survival. Embryonic cuticle formation is controlled by a signaling pathway involving the protease ALE1 and the receptor-like kinases GSO1 and GSO2. We show that a sulfated peptide, TWISTED SEED1 (TWS1) acts as a GSO1/GSO2 ligand. Cuticle surveillance depends on the action of ALE1 which, unlike TWS1 and GSO1/2, is not produced in the embryo but in the neighboring endosperm. Cleavage of an embryo-derived TWS1 precursor by ALE1 releases the active peptide, triggering GSO1/2-dependent cuticle reinforcement in the embryo. A bidirectional molecular dialogue between embryo and endosperm thus safeguards cuticle integrity prior to germination.

**One Sentence Summary:** Subtilase-mediated activation of the TWISTED SEED1 peptide provides spatial cues during embryo cuticle integrity monitoring.

## Main Text

In Angiosperms, seeds are composed of three genetically distinct compartments, the embryo and the endosperm resulting from the fertilization of the egg and central cell of the female gametophyte respectively, and the maternal seed coat. For the formation of viable seeds, development is coordinated by communication between the different seed compartments. In our work we have elucidated a bidirectional peptide-mediated signaling pathway between the embryo and the endosperm. This pathway regulates the deposition of the embryonic cuticle which forms an essential hydrophobic barrier separating the apoplasts of the embryo and endosperm. After germination, the cuticle - one of the critical innovations underlying the transition of plants from their original, aqueous environment to dry land - protects the seedling from catastrophic water loss (*1, 2*).

Formation of the embryonic cuticle has previously been shown to depend on two Receptor-Like Kinases (RLKs) GASSHO1/SCHENGEN3 (from here-on named GSO1) and GSO2, and on ALE1, a protease of the subtilase family (SBTs) (*2*–*5*). *gso1 gso2* and (to a lesser extent) *ale1* mutants produce a patchy, and highly permeable cuticle. These defects are not due to abnormal cuticle biosynthesis but instead are caused by defects in the spatial regulation of cuticle deposition (*2*). Mutant embryos also adhere to surrounding tissues causing a seed-twisting phenotype (*6*). Since SBTs have been implicated in the processing of peptide hormone precursors (*7, 8*), we hypothesized that ALE1 may be required for the biogenesis of the elusive inter-compartmental peptide signal required for GSO1/2-dependent cuticle deposition.

CASPARIAN STRIP INTEGRITY FACTORs (CIFs), a family of small sulfated signaling peptides, have recently been identified as ligands for GSO1 and GSO2 (*9*–*11*). CIF1 and CIF2 are involved in Casparian strip formation in the root endodermis (*11, 12*). The function of CIF3 and CIF4 is still unknown. To assess the potential role of CIF peptides in cuticle development, the quadruple mutant (*cif1 cif2 cif3 cif4)* was generated (Fig. S1A). Neither cuticle permeability nor seed twisting phenotypes were observed in this quadruple mutant (Fig. S1B-E). However, reduction (in the leaky *sgn2-1* allele (*12*)) or loss (in the *tpst-1* mutant (*13*)) of Tyrosyl-Protein Sulfotransferase (TPST) activity, results in seed-twisting and cuticle-permeability phenotypes resembling those observed in *ale1* mutants (Fig. 1A-D, Fig. S2 A-D). These data suggest that an as yet unidentified sulfated peptide may act as the ligand of GSO1/2 during seed development. Consistent with the hypothesis that TPST acts in the same pathway as GSO1 and GSO2, no difference was observed between the phenotype of *tpst-1 gso1-1 gso2-1* triple and *gso1-1 gso2-1* double mutants (Fig. S2E). In contrast, TPST and ALE1 appear to act synergistically, as a strong phenotype resembling that of *gso1 gso2* double mutants was observed in *tpst-1 ale1-4* double mutants (Fig. 1E-I) (Fig. S2F-J). This result supports the hypothesis that TPST and ALE1 act in parallel regarding their roles in embryonic cuticle formation, possibly through independent post-translational modifications that both contribute to the maturation of the hypothetical peptide signal.

**Fig. 1.**
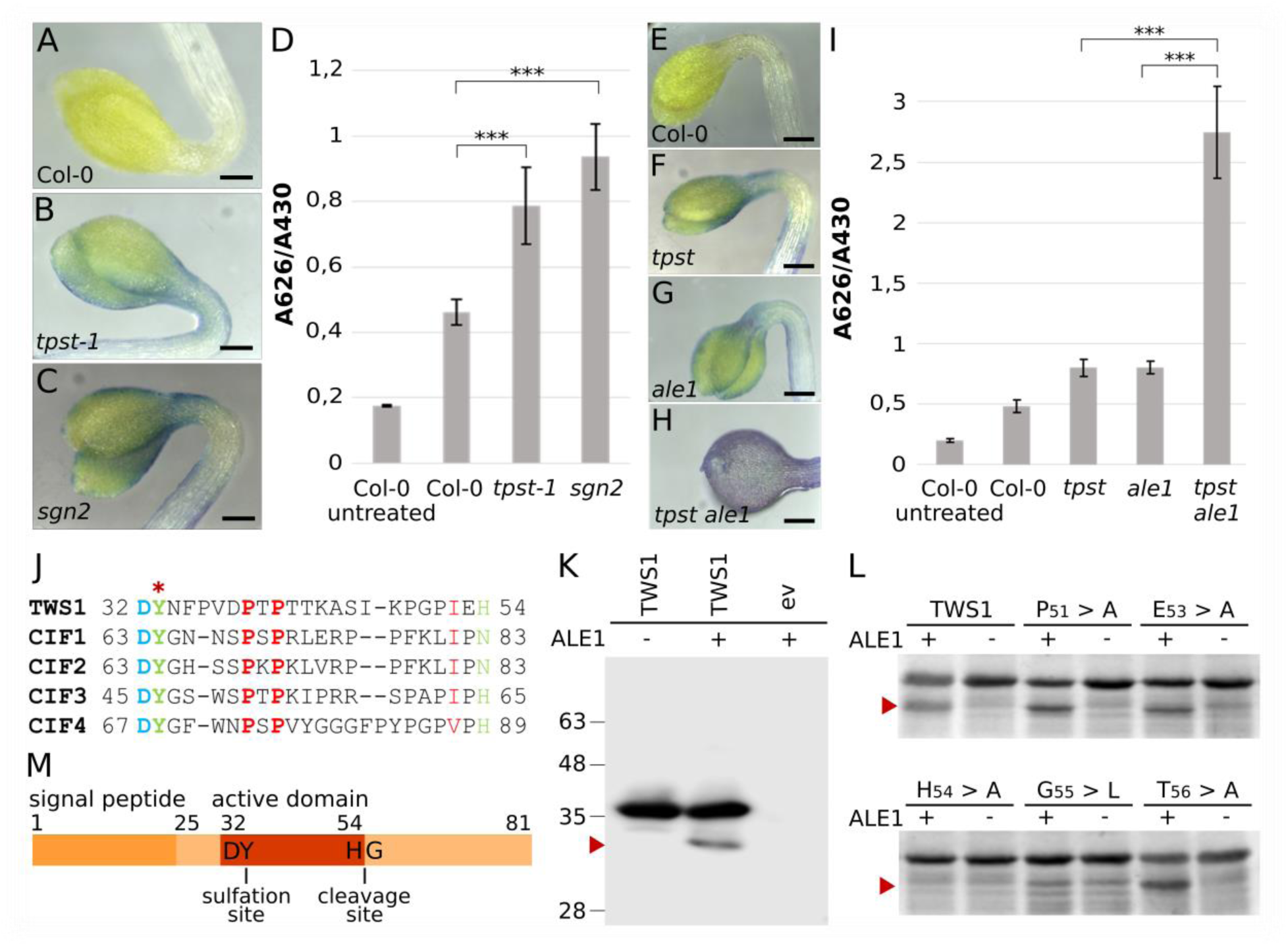
TPST and ALE1 are required for maturation of the TWS1 peptide. A-C and E-H) Toluidine blue tests on etiolated cotyledons. Scale bars = 200µm. D and I) Quantification of toluidine blue uptake by the aerial parts of young seedlings, normalized to chlorophyll content. N=6, ten seedlings per repetition. *** = statistical differences with one-way ANOVA followed by a *post-hoc* Scheffé multiple comparison test (P < 0,01) in D and I. J). The predicted TWS1 active peptide sequence and alignment with four other known GSO ligands (CIF1,CIF2, CIF3 and CIF4). The site of predicted sulfation is indicated with a red asterisk. K) Anti-GFP western blot of protein extracts from *N. benthamiana* leaves agro-infiltrated to express TWS1::GFP (TWS1) or the empty vector (-). Co-expression of ALE1::(HIS)6 or the empty-vector control are indicated by + and -, respectively. L) Coomassie-stained SDS-PAGE showing recombinant GST-TWS1 and the indicated site-directed mutants digested in vitro with (+) or without (-) ALE1-(His)6 purified from tobacco leaves. Arrows indicate specific cleavage products. M) The full length TWS1 precursor. Sulfation and ALE1 cleavage sites are indicated.

Identification of the peptide signal was facilitated by a recent study of TWISTED SEED1 (TWS1) (*14*), which despite not identifying TWS1 as a peptide precursor, reported a loss-of-function phenotype strikingly similar to that of *gso1 gso2* double mutants. Because existing alleles of *TWS1* are in the WS background, new CRISPR alleles (*tws1-3* to *tws1-10)* were generated in Col-0, and the phenotype of the resulting mutants was confirmed (Fig. S3). No additivity was observed when loss-of-function alleles of *TWS1* and of other pathway components (*GSO1, GSO2, TPST* and *ALE1*) were combined, providing genetic evidence for *TWS1* acting in the GSO signaling pathway (Fig. S4). Closer inspection of the TWS1 protein sequence revealed a region with limited similarity to CIF peptides including a DY motif which marks the N-terminus of the CIFs (Fig. 1J). The DY motif is the minimal motif required for tyrosine sulfation by TPST (*15*). Corroborating the functional importance of the putative peptide domain, the *tws1-6* allele (deletion of six codons in the putative peptide-encoding region) and the *tws1-5* allele (substitution of eight amino acids including the DY motif) both showed a total loss of function of the TWS1 protein (Fig. S3).

Next we tested whether TWS1 is a substrate of ALE1 by co-expression of ALE1:(His)6 and TWS1:GFP-(His)6 fusion proteins in tobacco (*N. benthamiana*) leaves. A specific TWS1 cleavage product was observed upon co-expression of ALE1 but not in the empty-vector control suggesting that TWS1 is processed by ALE1 *in planta* (Fig. 1K). Likewise, recombinant TWS1 expressed as GST-fusion in *E. coli* was cleaved by purified ALE1 *in vitro*. (Fig. 1L). Mass spectroscopy analysis of the TWS1 cleavage product purified from tobacco leaves showed that ALE1 cleaves TWS1 between His^54^ and Gly^55^ (Fig. S5). These residues are important for cleavage site selection, as ALE1-dependent processing was not observed when either His^54^ or Gly^55^ was substituted by site-directed mutagenesis (Fig. 1L). His^54^ corresponds to the C-terminal His or Asn of CIF peptides (Fig. 1J). The data thus suggest that ALE1-mediated processing of the TWS1 precursor marks the C-terminus of the TWS1 peptide. For CIF1 and CIF2, C-terminal processing is not required, as these peptides are located at the very end of their respective precursors. C-terminal processing could thus represent a specific mechanism of peptide activation operating in the developing seed but not in the root. A summary of TWS1 modifications is provided in Fig. 1M.

To test the biological activity of TWS1, the predicted peptide encompassing the conserved DY-motif at the N-terminus and the C-terminus as defined by the ALE1 cleavage site was custom-synthesized in tyrosine-sulfated form. As synthetic TWS1 cannot easily be applied to developing embryos, a root bioassay for CIF activity was used. In wild-type roots TWS1 induced ectopic endodermal lignification as previously observed for the CIF1 and CIF2 peptides (*12*). TWS1 activity was GSO1-dependent, suggesting that processed TWS1 peptide can replace CIF1 and CIF2 as a ligand for GSO1 during Casparian strip formation (Fig. 2A) (Fig. S6). Supporting this idea, TWS1 application complemented the phenotype of *cif1 cif2* mutants albeit with somewhat reduced activity compared to CIF2 (Fig. 2B, Fig. S7). TWS1 activity in this assay was strongly reduced, when the sulfate on the DY motif was missing (Fig. 2B). Versions of TWS1 in which Y^33^ was mutated to either F or T were only able to partially complement the mutant phenotype of *tws1-4*, consistent with a residual but weak activity for non-sulfated TWS1 *in vivo* (Fig S8). These results are consistent with the weak loss-of-function phenotype of the *tpst-1* mutant.

**Fig. 2.**
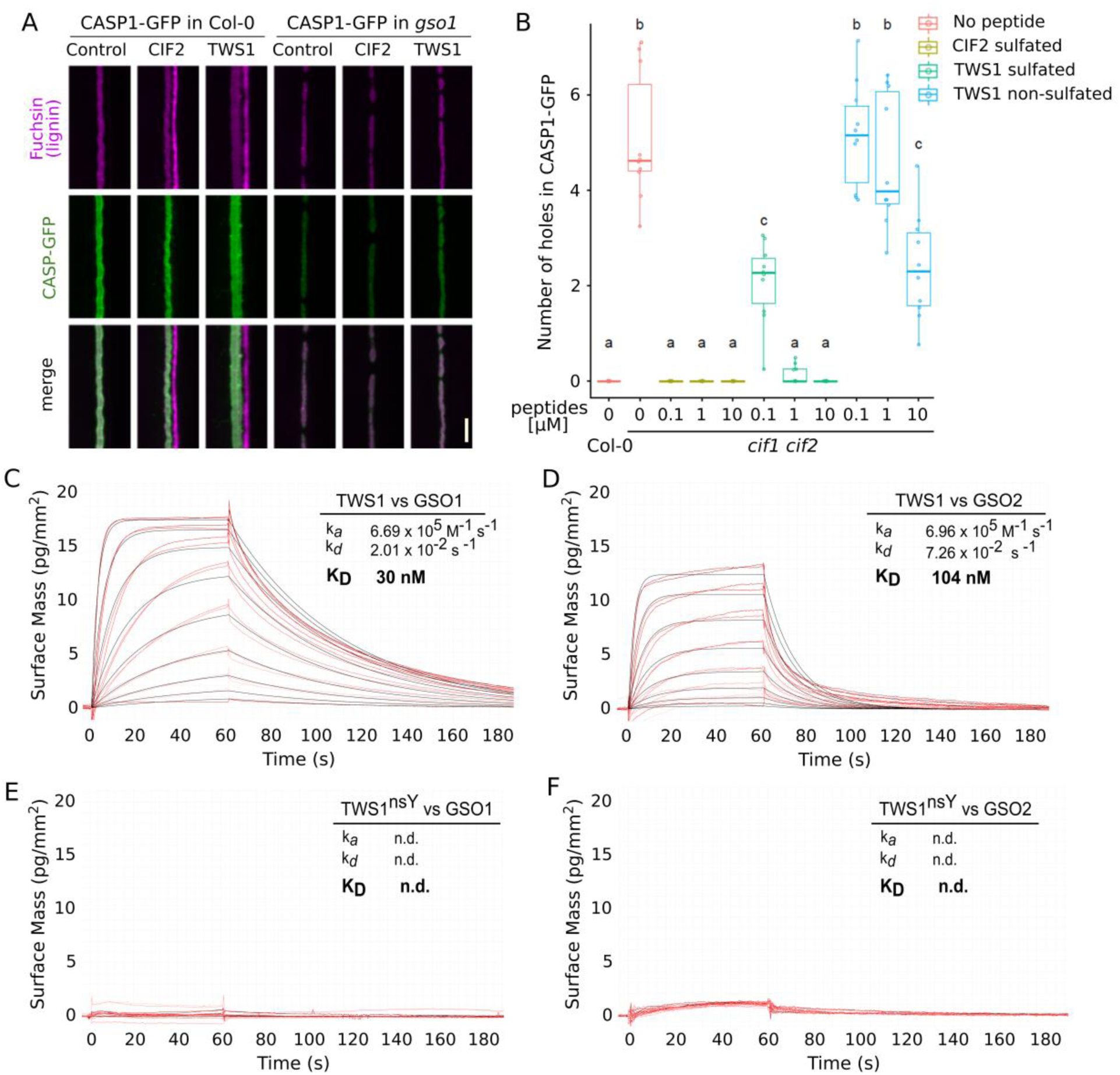
The TWS1 peptide is a functional GSO1/GSO2 ligand. A) Root over-lignification following treatment with the active CIF2 or TWS peptide in Col-0 and in the *gso1 (sgn3-3)* background. Lignin is stained in purple and CASP-GFP fusion protein, marking the Casparian strip domain, in green. Scale bar = 5µm B) Complementation of *cif1-2 cif2-2* Casparian strip integrity phenotype by peptide treatments. Number of holes in CASP-GFP counted after treatment with CIF2 sulphated peptide, TWS1 sulphated peptide, TWS1 non-sulphated peptide. N=10. a, b, and c correspond to a statistically supported classes supported by one-way ANOVA analysis followed by Tukey tests (P < 0,05). C-F) Grating-coupled interferometry (GCI)-derived binding kinetics. Shown are sensorgrams with raw data in red and their respective fits in black. *k*_a_, association rate constant; *k*_d_, dissociation rate constant; *K*_D_, dissociation constant. C) on the GSO1 extra-cellular domain in the presence of the sulphated TWS1 peptide. D) on the GSO2 extra-cellular domain in the presence of the sulphated TWS1 peptide. E) on the GSO1 extra-cellular domain in the presence of the non-sulphated TWS1 peptide. F) on the GSO2 extra-cellular domain in the presence of the non-sulphated TWS1 peptide.

To confirm TWS1 as a ligand of GSO1 and GSO2, the interaction of the synthetic peptide with the leucine-rich repeat (LRR) ectodomains of the receptors was analyzed in grating – coupled interferometry binding assays. GSO1 bound sulfated TWS1 with a K_*D*_ of ∼ 30 nM (Fig. 2C). The observed binding affinity is ∼10 fold lower compared to the CIF2 peptide (K_*D*_ = 2.5 nM) (Fig. S9), which is consistent with the reduced ability of TWS1 to complement the root phenotype of the *cif1 cif2* double mutant (Fig. 2B). Sulfated TWS1 also bound to the LRR domain of GSO2, albeit with slightly reduced affinity (K_*D*_ ∼ 100 nM) (Fig.2D). As previously shown for other CIF peptides (*10*), tyrosine sulfation was found to be critical for the interaction of TWS1 with GSO1 and GSO2 *in vitro* (Fig. 2E,F).

Taken together, our results suggest the sulfated TWS peptide as the missing link in the inter-compartmental signaling pathway for embryonic cuticle formation. The activities of ALE1 and TPST both contribute to the formation of the bioactive peptide (Fig. 1M) which is perceived by GSO1 and GSO2 to ensure appropriate cuticle deposition.

To understand how the elements of the signaling pathway cooperate to ensure the formation of a complete and functional cuticle, we analyzed their spatial organization in developing seeds. *In silico* data indicate that the *TPST* gene is expressed in all seed tissues (Fig. S10A) (*16, 17*). To investigate in which compartment of the seed TPST (which acts cell autonomously in the secretory pathway (*13*)) is explicitly required for TWS1 maturation, reciprocal crosses and complementation assays using tissue specific promoters were performed. Cuticle permeability defects were not observed when homozygous mutants were pollinated with wild-type pollen, confirming that defects are zygotic in origin (Fig. 3A-C). Cuticle defects of the *tpst-1* mutant were complemented when TPST was expressed under the *RPS5A* promoter that is active ubiquitously in the seed (*18*), and the *PIN1* promoter which drives expression in the embryo, but not in other seed tissues (Fig. S11). No complementation was observed for the endosperm-specific *RGP3* promoter (*19*), indicating that TPST activity is required for TWS1 sulfation specifically in the embryo to ensure cuticle integrity (Fig. 3D) (Fig. S12). Consistent with this observation, and with a previous report (*14*), the *TWS1* promoter was found to driving a strong and apparently uniform expression of TWS1 which was specific to the developing embryo from the early globular stage onwards (Fig. 3E) (Fig S13). A fluorescent reporter controlled by the *TPST* promoter (*12*) showed strong signal throughout the embryo proper at the onset of embryo cuticle establishment (globular stage) before becoming restricted to the root tip at later stages (Fig. S10). We conclude that the TWS1 peptide is both sulfated and secreted specifically in the embryo.

**Fig. 3.**
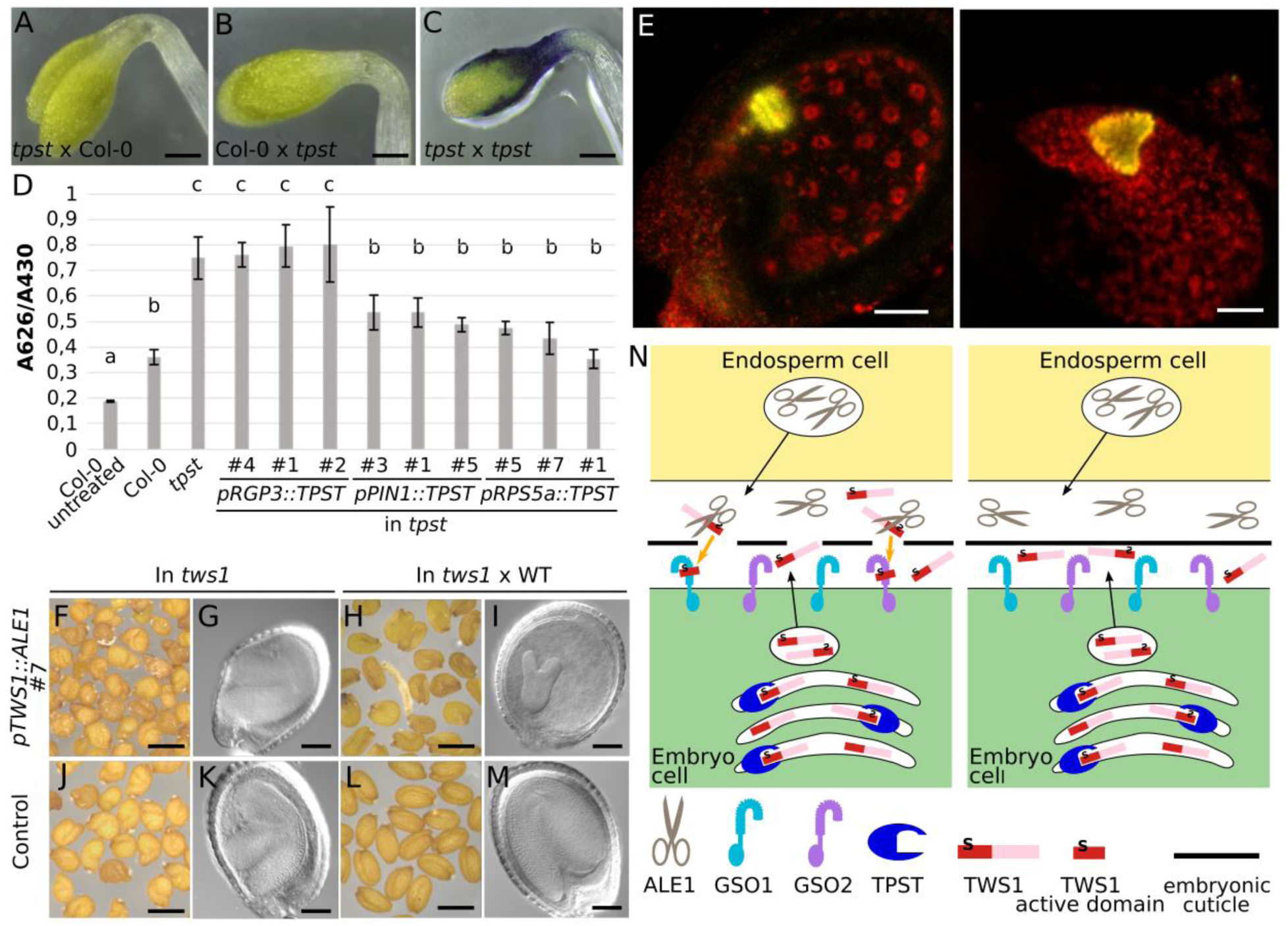
Spatial separation of *ALE1* and *TWS1* expression is critical for pathway function. A-C) F1 seedlings from reciprocal crosses stained with Toluidine blue. D) Complementation of *tpst-1* mutant with endosperm-specific expression of *TPST* (*pRGP3::TPST*), embryo-specific expression of *TPST (pPIN1:TPST)* and ubiquitous expression of *TPST (pRPS5a::TPST)* compared to *tpst-1* and Col-0. 3 independent lines were analysed. a, b, c = statistical differences with one-way ANOVA followed by a *post-hoc* Scheffé multiple comparison test (P < 0,01). E) Confocal images of *pTWS1::mCitrine::NLS-mCitrine* reporter lines, signal in yellow, autofluorescence in red. Scale bars = 50µm. F,G) Dry seeds and chloral hydrate cleared seeds (9 DAP) respectively from a line expressing *ALE1* in the embryo in the *tws1-4* background (*pTWS1::ALE1 line#7*). H,I) Seeds from crosses of Col-0 pollen onto *line#7*. J,K) self-fertilized *tws1-4* seeds as a control L,M) Seeds from a cross of Col-0 pollen on a *tws1-4* pistil as a control. Results for three further independent transgenic lines are shown in Figs. S17 and S18. N) Model for embryonic cuticle integrity monitoring.

However, the production of mature TWS1 requires a C-terminal cleavage event that we have shown to be mediated by ALE1. Importantly, *ALE1* is expressed only in the endosperm (*4, 20*), on the opposite side of the nascent cuticle to the GSO1 and GSO2 receptors, which are localized throughout the membranes of the embryo epidermal cells that produce the cuticle (Fig. S14-16) *(2)*. Our expression data thus support a model in which activation of the GSO signaling pathway depends on the migration of the TWS1 peptide precursor from the embryo to the endosperm, where it is cleaved and activated by ALE1 before diffusing back to the embryo to trigger GSO1/2-dependent cuticle deposition. Consequently, signaling is terminated as soon as the cuticle is complete, separating the subtilase from its substrate.

Genetic support for this model was obtained by ectopically expressing *ALE1* in the embryo, under the control of the *TWS1* promoter. Transformants could be obtained in *tws1* mutants, but not in the wild-type background. Furthermore, when *tws1* plants carrying the *pTWS1:ALE1* transgene were pollinated with wild-type pollen to introduce a functional *TWS1* allele into the zygotic compartments and thus induce colocalization of TWS1 precursors, ALE1 and the GSO receptors in the embryo, premature embryo arrest was observed (Fig. 3F-M) (Fig. S17, S18). The data indicate that co-expression of *TWS1* and *ALE1* in the embryo is highly detrimental to embryo development, likely due to constitutive embryonic activation of the *GSO1/GSO2* signaling pathway. Previous work has shown this pathway to activate the expression of stress-responsive genes in the seed context (*2*). We thus postulate that the spatial separation of the TWS1 precursor and the GSO receptors from the activating protease by the nascent cuticle is a prerequisite for the attenuation of signaling when a functional cuticle has been formed.

The proposed bidirectional signaling model allows efficient embryo cuticle integrity monitoring. The sulfated TWS1 precursor is produced by the embryo and secreted (probably after N-terminal cleavage of the pro-peptide) to the embryo apoplast. In the absence of an intact cuticular barrier, it can diffuse to the endosperm and undergo activation by ALE1 (and potentially other subtilases). Activated TWS1 peptide then leaks back through cuticle gaps to bind the GSO1 and GSO2 receptors and activate local gap repair (Figure 3N). When the cuticle is intact, proTWS1 peptides are confined to the embryo where they remain inactive.

Our results demonstrate a key role for a subtilase in providing spatial specificity to a bidirectional peptide signaling pathway. In contrast, the related CIF1, CIF2 and GSO1-dependent signaling pathway controlling Casparian strip integrity is uni-directional, negating the need for C-terminal cleavage-mediated peptide activation (*9, 11*). Consistent with this, CIF1 and CIF2 peptides lack the C-terminal extension present in the TWS1 precursor. Both pathway components and their spatial organization, differ markedly between the two systems, suggesting a potentially independent recruitment of the GSO receptors to different integrity monitoring functions within the plant.

## Acknowledgments

We would like to thank Loïc Lepiniec for providing the *tws1-1* and *tws1-2* seeds and Carlos Galvan Ampudia for the pPIN1::GFP seeds. We would like to acknowledge Audrey Creff and Sophy Chamot for their technical support and training. We would like to thank Alexis Lacroix, Patrice Bolland and Justin Berger for help with growing plants, Isabelle Desbouchages and Hervé Leyral for help with laboratory logistics, and Angélique Patole, Brigitte Martin Sempore, Cindy Vial and Stéphanie Maurin for administrative assistance. We also thank Berit Würtz and Jens Pfannstiel from the Core Facility Hohenheim for mass spectrometric analyses.

## Author contributions

G.I. led the study. G.I. and N.G. obtained funding for the study. G.I., N.G., A.Sc., M.H. T.W and A.St. supervised the work. N.M.D., S.R., S.F. and S.O. carried out the experiments. All authors were involved in the analysis of the results. G.I., A.Sc. and N.M.D. wrote the paper with input from all authors.

## Data and materials availability

All lines used in the study will be provided upon signature of an appropriate Material Transfer Agreement. All data is available in the main text or the supplementary materials.

## Supplementary Materials

Materials and Methods

Figures S1-S18

